# Interrelated effects of age and parenthood on whole-brain controllability: protective effects of parenthood in mothers

**DOI:** 10.1101/2022.07.13.499891

**Authors:** Hamidreza Jamalabadi, Tim Hahn, Nils R. Winter, Erfan Nozari, Jan Ernsting, Susanne Meinert, Elisabeth Leehr, Katharina Dohm, Jochen Bauer, Julia-Katharina Pfarr, Frederike Stein, Florian Thomas-Odenthal, Katharina Brosch, Marco Mauritz, Marius Gruber, Jonathan Repple, Tobias Kaufmann, Axel Krug, Igor Nenadić, Tilo Kircher, Udo Dannlowski, Birgit Derntl

**Author notes:** These authors contributed equally to this work (Hamidreza Jamalabadi, Tim Hahn). Corresponding authors: Prof. Dr. rer. nat. Hamidreza Jamalabadi, Department of Psychiatry and Psychotherapy, Philipps, University of Marburg, Germany, Rudolf-Bultmann Straße 8, D-35039 Marburg, / 5863827.

## Abstract

**Background:** Controllability is a measure of the brain’s ability to orchestrate neural activity which can be quantified in terms of properties of the brain’s network connectivity. Evidence from the literature suggests that aging can exert a general effect on whole-brain controllability. Mounting evidence, on the other hand, suggests that parenthood and motherhood in particular lead to long-lasting changes in brain architecture that effectively slow down brain aging. We hypothesize that parenthood might preserve brain controllability properties from aging.

**Methods:** In a sample of 814 healthy individuals (aged 33.9±12.7 years, 522 females), we estimate whole-brain controllability and compare the aging effects in subjects with vs. those without children. We use diffusion tensor imaging (DTI) to estimate the brain structural connectome. The level of brain control is then calculated from the connectomic properties of the brain structure. Specifically, we measure the network control over many low-energy state transitions (average controllability) and the network control over difficult-to-reach states (modal controllability).

**Results and conclusion:** In nulliparous females, whole-brain average controllability increases, and modal controllability decreases with age, a trend that we do not observe in parous females. Statistical comparison of the controllability metrics shows that modal controllability is higher and average controllability is lower in parous females compared to nulliparous females. In men, we observed the same trend, but the difference between nulliparous and parous males do not reach statistical significance. Our results provide strong evidence that parenthood contradicts aging effects on brain controllability and the effect is stronger in mothers.

## 1. Introduction

Across species, the female brain experiences dynamic changes over the course of pregnancy and the postpartum period which are thought to support maternal adaptations (Duarte-Guterman, Leuner, & Galea, 2019; Hoekzema et al., 2017; Hoekzema et al., 2020). While some of these alterations are short-lived, accumulating evidence suggests long lasting structural changes to be traceable not only years (Hoekzema et al., 2017; Hoekzema et al., 2020) but also decades after childbirth in females (A.-M. G. de Lange et al., 2019). Specifically, a higher number of children was associated with less grey matter brain aging in limbic and striatal regions as well as the hippocampus (A. M. G. de Lange et al., 2020). Similarly, investigating white matter brain age, Voldsbekk and colleagues (Voldsbekk et al., 2021) reported that a higher number of childbirths was related to lower brain age in global white matter and in specific tracts, with the corpus callosum contributing uniquely to this association. Besides highlighting the high plasticity of maternal brain architecture, these structural changes are also associated with diminished aging effects in a wide range of cognitive abilities including memory, learning, and general brain health (Richard et al., 2018). On a cellular level, motherhood was associated with significantly elongated telomeres, pointing to slower cellular aging (Barha & Galea, 2017). Taken together, there is converging evidence that transition to motherhood involves vast physiological (e.g., pregnancy) and environmental changes (e.g., parenting), including changes to the brain structure, but the contribution and interaction of these changes and their impact on the maternal brain have remained unclear.

Physiological and environmental changes, though, are experienced by both parents. Neurobiological literature on nonhuman fathers indicates that becoming a father involves a major neurohormonal reorganization that prepares for the expression of adequate caregiving across mammalian species (for review see Swain, Dayton, Kim, Tolman, & Volling, 2014). In humans, recently, Ning et al. (Ning et al., 2020) reported better visual memory and faster response times in mothers and fathers compared to non-parents, with protective effects on cognitive function being larger in males than females. In terms of brain age, the authors further showed a significant decline in mothers and fathers compared to non-parents, though here the effect is stronger in mothers. Furthermore, Orchard and colleagues (Orchard et al., 2020) reported significant differences in cortical thickness between parents and non-parents: For mothers, a significant positive association between number of children and cortical thickness of the right parahippocampal gyrus as well as improved verbal memory performance emerged. Investigating a potentially persistent effect of parenthood on resting-state functional connectivity in older adults, the same group (Orchard et al., 2021) observed, only in females, a significant association between decreases in functional connectivity and number of children parented. More specifically, increased segregation between networks, decreased connectivity between hemispheres, and decreased connectivity between anterior and posterior regions were reported for mothers only. The authors conclude that their findings suggest a beneficial effect of motherhood for brain function in late life.

Taken together, these findings suggests that motherhood - and potentially fatherhood - might affect whole brain structural properties in a way that preserves the brain from aging. While a detailed mechanistic model linking the structural brain properties to the cognitive abilities and functional dynamics is still lacking, network control theory has shown to be a valuable tool enabling systematic analysis of those relationships. Network control quantifies the dynamic behavior of brain activity in response to the external, as well as internal, perturbation. Within the framework of network control, controllability centralities refer to the properties of single nodes to steer the functional network dynamics (Jamalabadi et al., 2021). In particular, average and modal controllability have been suggested to quantify the brain’s ability to execute easy and difficult state transitions, respectively, and have been shown to be sensitive enough to relate to a wide range of brain cognitive and functional properties (Betzel, Gu, Medaglia, Pasqualetti, & Bassett, 2016; Gu et al., 2015; Hahn et al., 2021; Tang & Bassett, 2018) as well of normal human aging (Tang et al., 2017).

In this study, we hypothesized that parenthood might affect age-related whole brain controllability, in line with recent reports on brain aging (A.-M. G. de Lange et al., 2019; A. M. G. de Lange et al., 2020; Ning et al., 2020). Based on recent reports linking motherhood with white matter brain age (Voldsbekk et al., 2021) and the heterogenous results on the paternal brain (Ning et al., 2020; Orchard et al., 2021; Orchard et al., 2020 vs), we further expected those effects to be stronger in mothers.

## 2. Methods

### Sample

Participants were healthy individuals as part of the Marburg-Münster Affective Disorders Cohort Study (MACS) (Kircher et al., 2019) and were recruited at two different sites (Marburg & Münster, Germany) (Vogelbacher et al., 2018). Participants ranging in age from 18 to 65 years were recruited through flyer and newspaper advertisements. All experiments were performed in accordance with the ethical guidelines and regulations and all participants gave written informed consent prior to examination. Exclusion criteria comprised the presence of any lifetime mental disorder, neurological abnormalities, history of seizures, head trauma or unconsciousness, major medical conditions (e.g., cancer, unstable diabetes, epilepsy etc.), current pregnancy, hypothyroidism without adequate medication, claustrophobia, color blindness, and general MRI contraindications (e.g., metallic objects in the body). In total we had access to data from 814 healthy participants (522 females, 292 males) including 369 parents (167 mothers, 202 fathers). We compared education years and verbal intelligence test scores (MWTB; Lehrl, 2005) across subsamples (nulliparous females, nulliparous males, parous females, parous males), and do not find any significant group differences. However, in terms of age, the nulliparous and parous groups differed significantly, with the parous females and males being on average older than the nulliparous females and males. Therefore, we selected all nulliparous females and males aged older than 30 years to serve as an additional control group without any significant age difference between parous vs. nulliparous individuals. Table 1 summarizes the demographic information.

**Table 1:** Demographic information of the participants included in the current study including age, years of education and premorbid verbal intelligence (Mehrfachwahl-Wortschatz-Intelligenztest B, MWTB). Since the nulliparous subsamples have a significant lower average age, we replicate all analysis in two groups where we first exclude and then include the samples below 30 years old.

|  | Parous<br>Females | Parous<br>Males | Nulliparous<br>Females<br>(> 30 years) | Nulliparous<br>Males<br>(> 30 years) | Nulliparous<br>Females (all) | Nulliparous<br>Males (all) |
| --- | --- | --- | --- | --- | --- | --- |
| **No.** | 167 | 202 | 67 | 50 | 355 | 90 |
| **Age (mean ± std)** | 47.6 ± 9.3 | 47.1 ± 9.4 | 41 ± 9.2 | 39.5 ± 9.6 | 27.2 ± 8.5 | 28.4 ± 8.5 |
| **Years of education**<br>**(mean ± std)** | 13.3 ± 2.9 | 14.4 ± 3.2 | 14.76 ± 2.9 | 14.6 ± 3.4 | 14.1 ± 2.2 | 14.0 ± 2.4 |
| **MWTB raw score**<br>**(mean ± std)** | 31.6 ± 2.8 | 31.7 ± 3.2 | 32.0 ± 2.7 | 31.3 ± 3.5 | 30.7 ± 2.9 | 30.3 ± 3.2 |

### Imaging data acquisition

In the MACS, two MRI scanners were used for data acquisition located at the Departments of Psychiatry at the University of Marburg and the University of Münster, both Germany, with different hardware and software configurations. Both T1 and DTI data were acquired using a 3T whole body MRI scanner (Marburg: Tim Trio, 12-channel head matrix Rx-coil, Siemens, Erlangen, Germany; Münster: Prisma, 20-channel head matrix Rx-coil, Siemens, Erlangen, Germany). A GRAPPA acceleration factor of two was employed. For DTI imaging, 56 axial slices, 2.5 mm thick with no gap, were acquired with an isotropic voxel size of 2.5 mm^3^ (TE = 90 ms, TR = 7300 ms). Five non-DW images (b0=0) and 2 x 30 DW images with a b-value of 1000 sec/mm^2^ were acquired. Imaging pulse sequence parameters were standardized across both sites to the extent permitted by each platform. For a description of MRI quality control procedures see (Vogelbacher et al., 2018). The body coil at the Marburg scanner was replaced during the study. Therefore, two variables modeling three scanner sites (Marburg old body coil, Marburg new body coil and Münster) were used as covariate for all statistical analyses

### Preprocessing of diffusion-weighted images and connectome reconstruction

Diffusion-weighted images (DWI) were realigned and corrected for eddy currents and susceptibility distortions using FSL’s eddy (Version 6.0.1). Diffusion tensor imaging models the measured signal of a voxel by a single tensor describing the diffusion signal as one preferred diffusion direction per voxel. Connectomes were reconstructed using the CATO toolbox (See Supplement 4) (S. C. de Lange & van den Heuvel, 2021). CATO uses the informed RESTORE algorithm that estimates the tensor while identifying and removing outliers during the fitting, thereby reducing the impact of physiological noise artifacts on the DTI modeling. Based on the diffusion profiles, white matter pathways were reconstructed using deterministic tractography. To this end, eight seeds were started per voxel, and for each seed, a tractography streamline was constructed by following the main diffusion direction from voxel to voxel. Stop criteria included reaching a voxel with a fractional anisotropy < 0.1, making a sharp turn of >45°, reaching a gray matter voxel, or exiting the brain mask.

For each subject an anatomical brain network was reconstructed, consisting of 114 cortical areas of a subdivision of the FreeSurfer’s Desikan–Killiany atlas (Cammoun et al., 2012; Hagmann et al., 2008), and the reconstructed streamlines between these areas. Given the poorer DWI signal-to-noise ratio in subcortical regions and the dominant effect of subcortical regions on network properties, we decided to use a subdivision of this atlas containing only cortical regions. White matter connections were reconstructed using deterministic streamline tractography, based on the Fiber Assignment by Continuous Tracking (FACT) algorithm (Mori, 2002). Network connections were included when two nodes (i.e., brain regions) were connected by at least three tractography streamlines (de Reus & van den Heuvel, 2013). For each participant, the network information was stored in a structural connectivity matrix, with rows and columns reflecting cortical brain regions, and matrix entries representing graph edges. Edges were only described by their presence or absence to create unweighted graphs.

### Network Controllability Analysis

Following the mainstream in the literature of brain controllability analysis (Karrer et al., 2020), we assume a noise-free linear time-invariant model (for a more detailed introduction, see (Gu et al., 2015)):

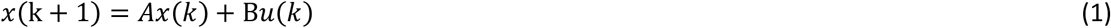

where the vector *x* represents the temporal activity of 114 brain regions, *A_114×114_* is the non-negative unweighted adjacency matrix representing the structural connectivity (see *Imaging Data Preprocessing* above for details), the scalar value *u*(*t*) represents the input (source of energy or activation) to the system, and the binary-valued vector *B* encodes the brain regions that distribute the input energy across the system.

For this system, **Modal Controllability** (*MC*) of node *i* is defined as 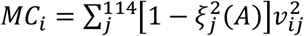 where *ξ_j_* and *V_ij_* are the eigenvalues and eigenvectors of *A*, respectively. Whole-brain modal controllability 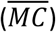 is then defined as the average of modal controllability over all nodes. Using the symmetry of *A* (and hence the orthonormality of its eigenvectors), this simplifies to:

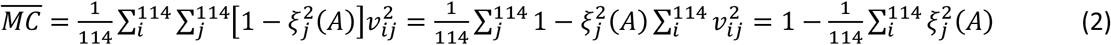

Note that 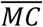 is now a function of the eigenvalues only. Thus, given that the eigenvalues determine the decay rate of the response to any arbitrary input *u*(*t*) (Kailath, 1980), equation (3) implies that whole-brain MC is negatively proportional to the duration of the output signal and therefore to the output power over a fixed interval of time.

Also, for the system defined in (1), **Average Controllability** (*AC*) measures the energy content of the system’s impulse response and is defined as *Tr*(*W_c_*) where 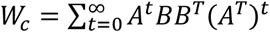 is the controllability Gramian of the system and *Tr*(.) denotes the trace of a matrix. Note that for a linear time-invariant system of this form, the impulse response fully determines the system’s response to any (not necessarily impulse) inputs (Oppenheim, Willsky, & Nawab, 1996). Average controllability for a single node *i* (i.e., when B includes only one nonzero element equal to one) simplifies to:

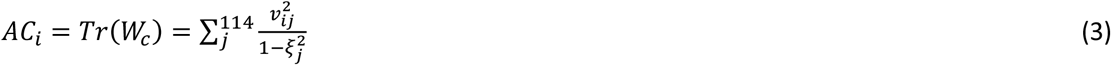

Thus, whole-brain (i.e., mean) average controllability over all nodes becomes

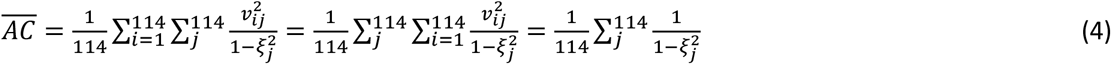

Note the similarity between equation (2) and equation (4), where the effects of the eigenvectors disappear when considering the whole-brain (i.e., averaging over all regions) controllability. Also, note the opposite dependence on the system modes where larger eigenvalues increase the whole-brain AC and thus, opposite to the whole-brain MC, is associated with longer output response and higher output power.

### Statistical Analyses

Given that our data are not balanced with respect to sex and parenthood, we conducted three consecutive stepwise analyses to empirically test our hypotheses without losing data to age-sex-parenthood balancing: First we trained a linear model to estimate AC based on age for two samples of nulliparous and parous females. Next, we tested if the age-related beta values are statistically moderated by parenthood (Cohen, Cohen, West, & Aiken, 2013) by training a full model including parenthood and age-parenthood interaction. We repeated the same procedure for MC and males. In all analyses, we included study site and the total number of present edges as co-variates. All statistical analyses were carried out using MATLAB.

## 3. Results

To empirically test our main hypothesis, we trained a linear model on subjects above 30 years old (224 women, aged 46.8 ± 9.1 years; 138 men, aged 44.8 ± 9.9 years). This decision was motivated by the characteristics of our data where most parents (94%) were aged 30 years or older (see Supplementary Figure S1). To ensure that our results are not confounded by this selection, we replicated all our analyses with different age intervals (see Supplementary Table S13 for details).

### Females

In nulliparous females aged over 30 years (67 females, aged 41.9 ± 9.2), whole-brain average controllability 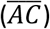 is positively (*β* = 0.29, *p* = 0.01) and whole-brain modal controllability 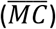 is negatively associated with age (*β* = −0.26, *p* = 0.03) (see full models in Supplementary Tables S1 and S2). Performing this test in mothers (157 mothers, aged 48.8 ± 8.2) however shows reversed but statistically nonsignificant effects (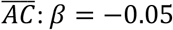, *p* = 0.51; 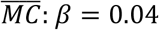, *p* = 0.62; see full models in Supplementary Tables S3 and S4). Directly comparing the beta values (see Figure 1A, Supplementary Figure S2A, Supplementary Figure S4) indicates a significant difference for both parameters: 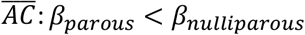, *P* =0.01; 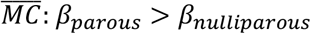, *P* = 0.02.

**Figure 1:**
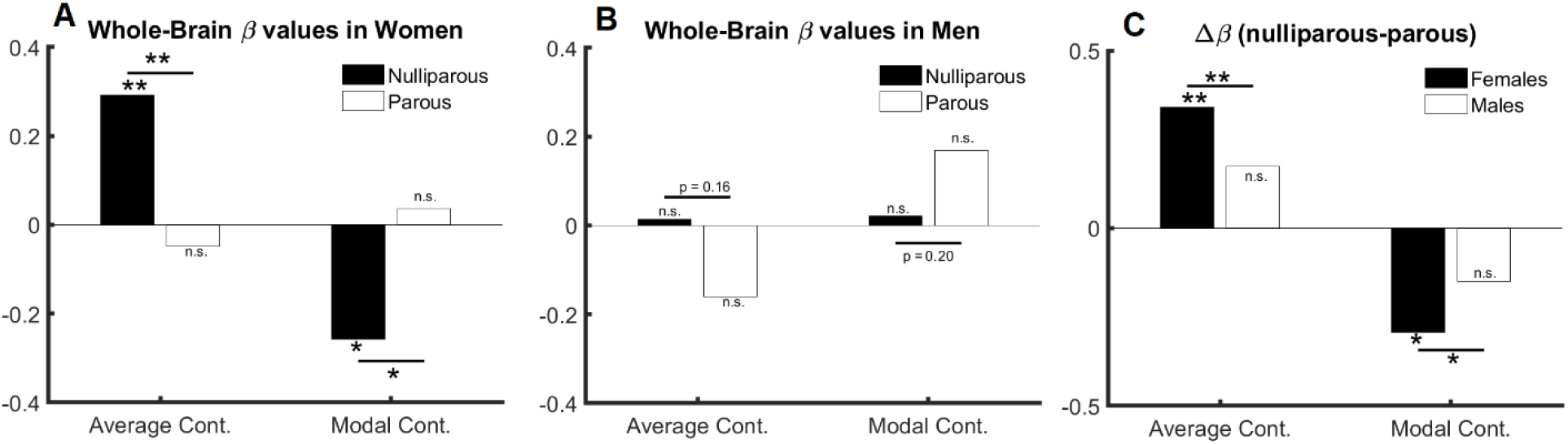
The relation between age, sex, and controllability (age > 30). (A) age and parenthood effect in females: average and modal controllability change with age only in nulliparous females. (B) age and parenthood effect in males: average and modal controllability change do not change with age in men. (C) Parenthood effect in males in comparison with females: although the change in beta values do not reach significance in males, they follow numerically the same pattern as in females.

The significant difference between age-related beta values in parous versus nulliparous females suggests an interaction between age and parenthood. We, therefore, trained an additional model where age, parenthood and age-parenthood interactions are included as independent variables in the linear model. The results indicate a significant interaction between age and parenthood in both controllability measures: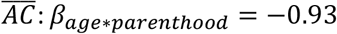, *p* = 0.008; 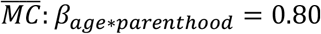, *p* = 0.02 (see Supplementary Tables S5 and S6 for full models).

### Males

Having established the effect of age and parenthood on whole-brain controllability in females, we asked whether the same effects would be observable in males where we trained the models from Supplementary Tables S5 and S6 on data from male subjects. Toward this end, none of our tests resulted in statistically significant effects for age, parenthood, or their interaction (all p > 0.19, see full results in Supplementary Tables S7 and S8). However, the changes in beta values follow the same structure as in females (see Figures 1B, Supplementary Figure S4).

We therefore performed three additional analyses: First, we pooled the data from females and males and trained the previous model without including sex as a covariate. We observe a significant interaction between age and parenthood for both parameters of brain controllability: 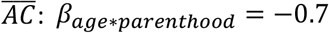, *P* = 0.01, 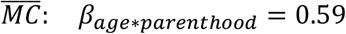, *p* = 0.03 (see Supplementary Tables S9 and S10 for full models).). Second, we compared the beta values for the models based on parous and nulliparous males, which indicates a smaller (and nonsignificant) difference in the same direction as in females: 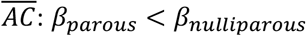, *P* = 0.16;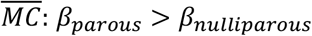, *P* = 0.20. Third, we extended the model to include sex as an additive co-variate. This model revealed age * parenthood interaction effects comparable to the model without including sex as a covariate (see full models in Supplementary Tables S11 and S12. Please see Figure 1 for effects of age, sex, and parenthood on average and modal controllability.

### Regional specificity

Finally, we tested if the compensatory effects of parenthood on age-related changes are specific to certain brain regions, excluding again the young nulliparous subset. Toward this end, we trained 114 models corresponding to 114 cortical regions of the brain and compared the corresponding beta values for age, parenthood, and age-parenthood interaction on the data from females (see figure 2; for data on males and mixed-sex analysis please see supplementary figures S2 and S3). The strong linear trend between the beta-values (*R*^2^ > 0.6) shows a strong effect of parenthood on controllability values that contracts that of age. Importantly, although average controllability increases with age on average, average controllability decreases with age in many regions. Among those regions, the largest effects belong to the right postcentral and left precentral regions. Related to this analysis, we further trained models in four groups of nulliparous and parous females and males and observes that compared to the nulliparous subjects, a large number of brain regions does not show age-related increase in average controllability (p=0.03 in females and p = 0.15 in males).

**Figure 2:**
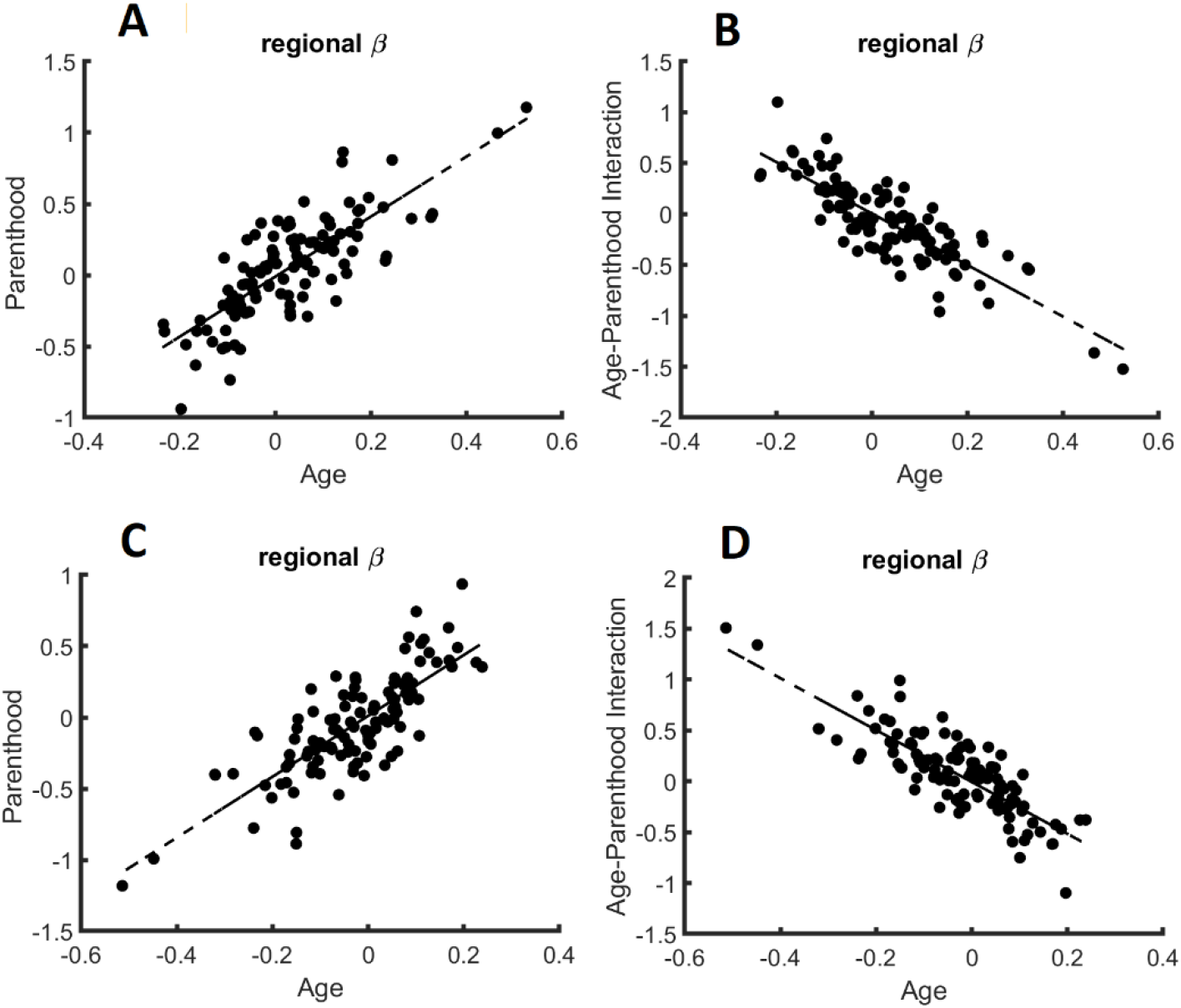
The relation of regional age and parenthood beta values in the linear models that predict whole-brain controllability in females. Similar to the whole-brain models of controllability, the beta values for age and parenthood have the same signs for models of average (A) and modal (C) controllability while age and age-parenthood have opposite signs in models of (B) average and (D) modal controllability.

## 4. Discussion

In this work, we estimated whole-brain controllability properties based on structural connectome data, that quantify the brain’s ability to perform state transitions (Karrer et al., 2020), in a group of healthy females and males, who were either parents or non-parents. Controllability, as we discussed in this paper, is a centrality metric that relates the activity of brain regions to their ability to steer the whole brain dynamics. On the node level, high controllability values relate to stronger and lower values indicate smaller ability to affect brain dynamics. On the whole brain level, the controllability metrics specifically code for the rate of signal decay in the brain following external or internal perturbation. Here, we aimed to investigate changes in brain controllability due to age and parenthood and to compare them between sexes. We observed that parenthood affects brain controllability stronger in females than males although the direction of change seemed to be similar. Specifically, our results point to compensating effects of parenthood on age-related changes in brain controllability, suggesting less brain aging in mothers. Our findings extend the previously reported effects of motherhood on brain aging using gray (Buchheim et al., 2006; A.-M. G. de Lange et al., 2019; A. M. G. de Lange et al., 2020) and white matter (Voldsbekk et al., 2021), adding a new dimension in terms of the topological complexity of these changes. Our results warrant the conclusion that the supporting effects of parenthood might be optimized topologically to support certain neuronal state transitions which we quantified here in terms of controllability metrics.

The effect of aging on brain controllability as a structural predictor of the ease by which the brain can switch from one dynamical state to another has also been investigated before using diffusion tensor imaging (Tang et al., 2017). In a neurodevelopmental cohort, authors showed an increase in average controllability and modal controllability as children age, suggesting that brain networks develop explicitly to maximize controllability during adolescence, i.e. adapting white matter connectivity to increase the ability to flexibly move between diverse brain states (Chai et al., 2017). However, data on how aging affects brain controllability in the adult brain are not available. While we observed an increase in average controllability and a decrease in modal controllability, indicating that brain dynamics shift with age favoring switches between easy to reach brain states, i.e., average controllability (as opposed to difficult to reach brain states reflected by modal controllability), this needs to be further investigated and replicated by future studies.

Despite previous data indicating brain malleability due to fatherhood, the interaction of age-by-parenthood on brain controllability measures did not reach significance in males in our study. Whether this observation mirrors a null effect or whether our sample was too small to detect the relatively small effect of parenthood on male brain controllability measures however requires further investigation. We however notice that, similar to our study, Orchard and colleagues (Orchard et al., 2021) did not report any significant effect of parity on functional connectivity components in fathers, only in mothers and that the male brain shows generally more stable controllability across the life-span in our study. Taken together, these findings might motivate the conclusion that the male brain is less plastic with respect to the effects of parenthood. Besides obvious biological differences between females and males (i.e., pregnancy, lactation), Orchard and colleagues cautiously speculate about the impact of “traditional” caregiving arrangements, which may impact the investigated functional connectivity patterns but were not collected in their study. Notably, parenthood is associated with several changes in life-style factors, including reduced alcohol and tobacco consumption (Kravdal, 1995). Moreover, children can connect parents to more social and community activities (Furstenberg, 2005), can provide emotional and social support (Ross & Mirowsky, 2002) and altogether may contribute to the lower mortality risk in parents in both sexes (Modig, Talbäck, Torssander, & Ahlbom, 2017). Whether these effects are again boosted by experiences of becoming aunt/uncle, grandparent, godparent, or any other significant affiliation with children, is unknown but possible. It may also be a question of attachment and quality of the relationship. Here, previous data from one study indicate that changes in regional brain volume are associated with mother-to-infant attachment not only in the early postpartum phase but also six years later (Hoekzema et al., 2017; Martínez-García et al., 2021). And results from (Abraham et al., 2014) indicate that parent-infant synchrony is associated with amygdala activation in mothers and superior temporal sulcus activation in primary caretaking fathers. However, studies relying on big data bases – including our study - rarely have information on the quality of attachment and quantity/quality of care taking which needs to be addressed in future studies.

This study has some imitations. Most importantly, it is an observational cross-sectional study, and as such one cannot causally state that parenthood is leading to beneficial effects on healthy brain aging based on our results. It could also be that, e.g., those who have poor underlying health have fewer opportunities to have children. Longitudinal studies including females and males before/during pregnancy, postpartum and child rearing phase up to midlife and older age are required to enable an insight on the causal effects of parenthood on human brain and behavior. Further, we did not have information on several confounding factors including parental sensitivity and number of children. This may add worthwhile information since e.g. in males, the association between parity and regional brain age is quadratic, i.e. a “U-shaped” function, while in females this association seems to be more linear (Ning et al., 2020). Authors speculate that these differences may be linked to hormonal variations due to pregnancy and lactation which needs to be further investigated. While we report long-term effects of parenthood on brain controllability measures, further studies are needed to better clarify the underlying mechanisms of how parenthood shapes brain controllability properties which, in addition to white matter, might also engage gray matter adaptations. Previous studies have shown that, in addition to white matter, age-related changes and protective effects of parenthood are also identifiable in grey matter and its related processes seem to be partially independent from that governing white matter adaptations (Voldsbekk et al., 2021). Relatedly, we showed that controllability properties are strongly dependent on the interactive levels of nodal gray matter volume and white matter tracts (Jamalabadi et al., 2021). Taken together, these studies suggest that an integral view of the parental brain should involve simultaneous analysis of gray as well as white matter.

## 5. Conclusion

We studied the interrelation of age, sex, and parenthood on brain controllability properties and observed a significant association between parenthood and brain controllability in females. In terms of structural brain connectivity, measured via diffusion tensor imaging, this is the first study to investigate effects of parenthood and age in adult females and males. Our results support the notion that parenthood preserves brain structure from aging and provides evidence for beneficial effects of parenthood particularly in mothers.

## Acknowledgment

The authors would like to thank Teresa Luther for constructive feedbacks regarding the results presented in this paper.

## Availability of data

The data that support the findings of this study are from https://for2107.de/. Restrictions apply to the availability of these data, which were used under license for this study. Data are available from authors with the permission of the https://for2107.de/consortium.

## Author contribution statement

### Conceptualization

HJ, TH, TK, BD. Methodology and Validation: HJ, TH, BD, EN, NRW, MM, MG, JR.

### Data Curation

TK, UD, IN, SM, EL, KD, JB, JKP, FS, FTO, KB. Writing-Original Draft: HJ, BD, Writing-Review: all.

## Supplementary Information

**Figure S1:**
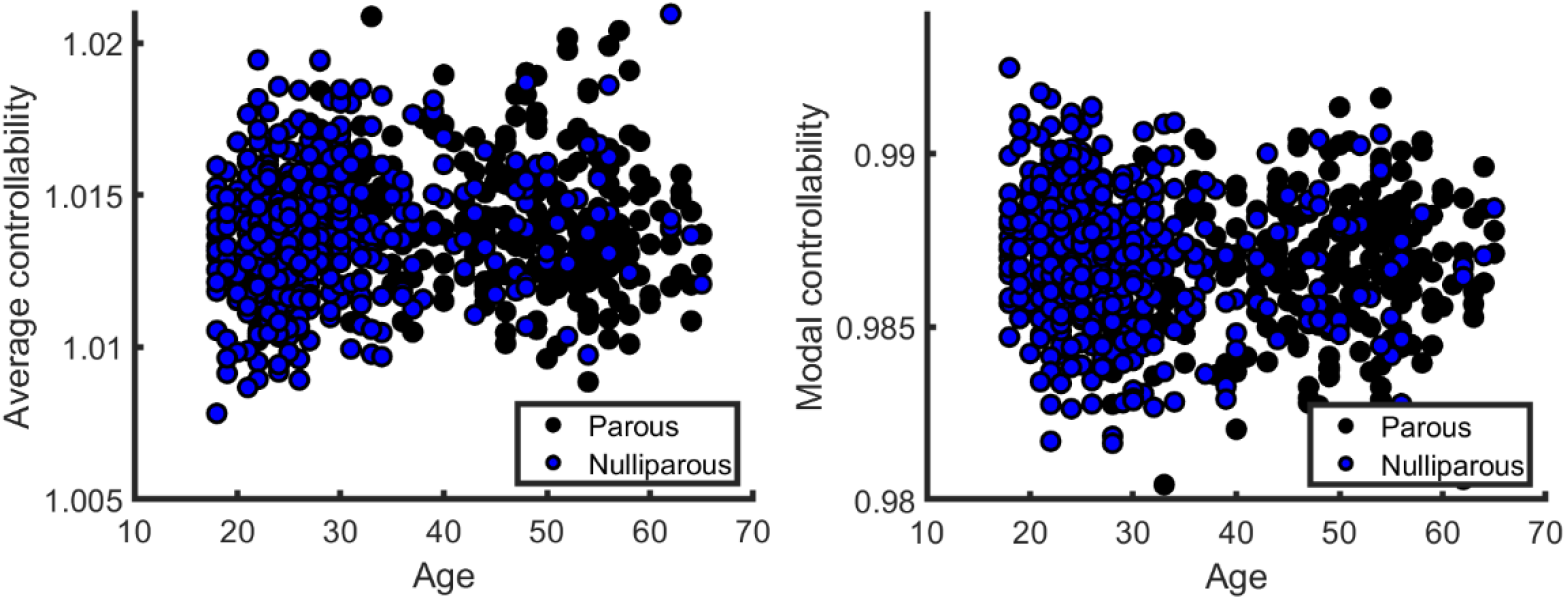
Distribution of average and modal controllability values for all data in this study.

**Figure S2.**
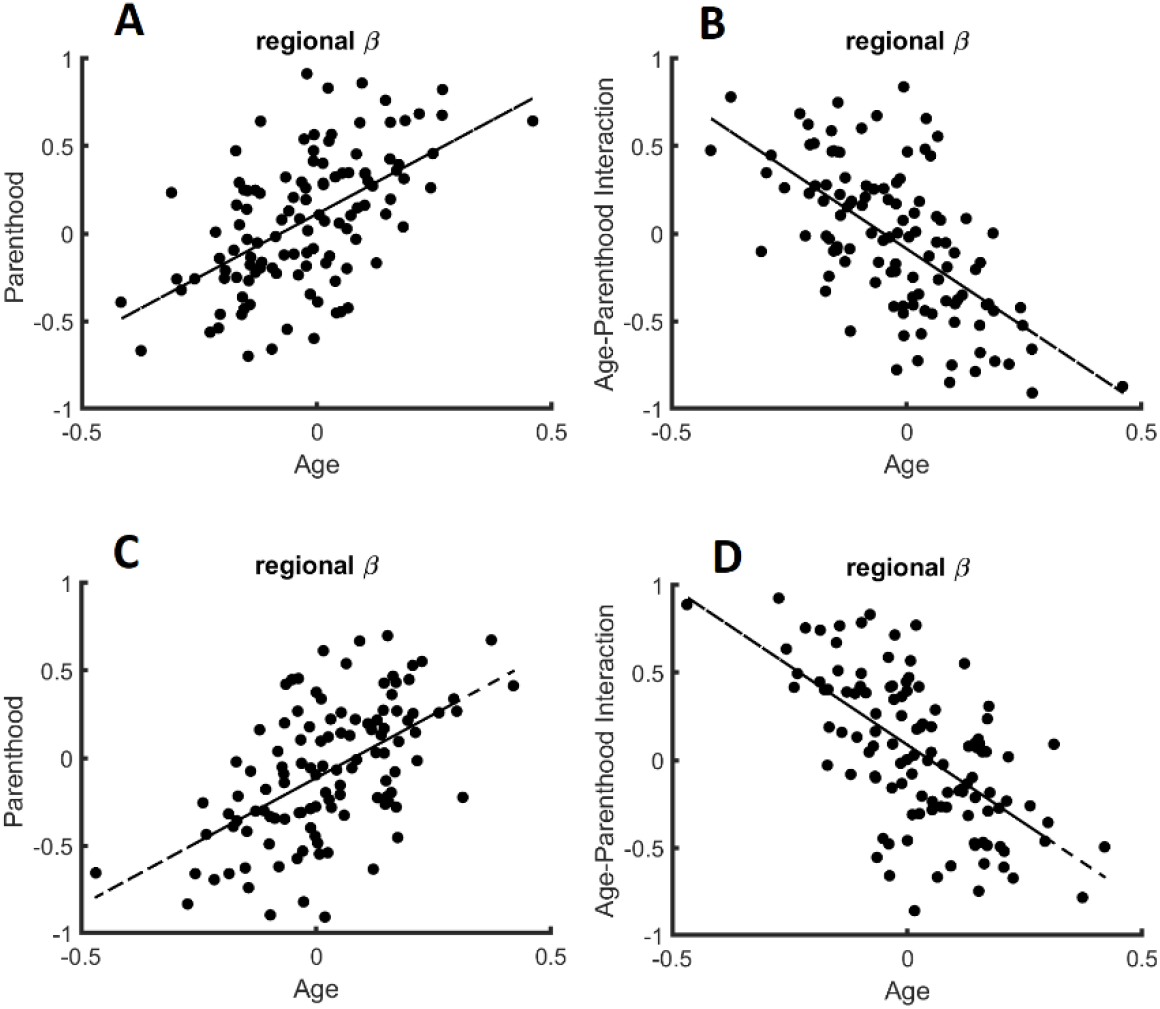
The relation of regional age and parenthood beta values in the linear models that predict whole-brain controllability in males. Similar to the whole-brain models of controllability, the beta values for age and parenthood have the same signs for models of average (A) and modal (C) controllability while age and age-parenthood have opposite signs in models of (B) average and (D) modal controllability.

**Figure S3.**
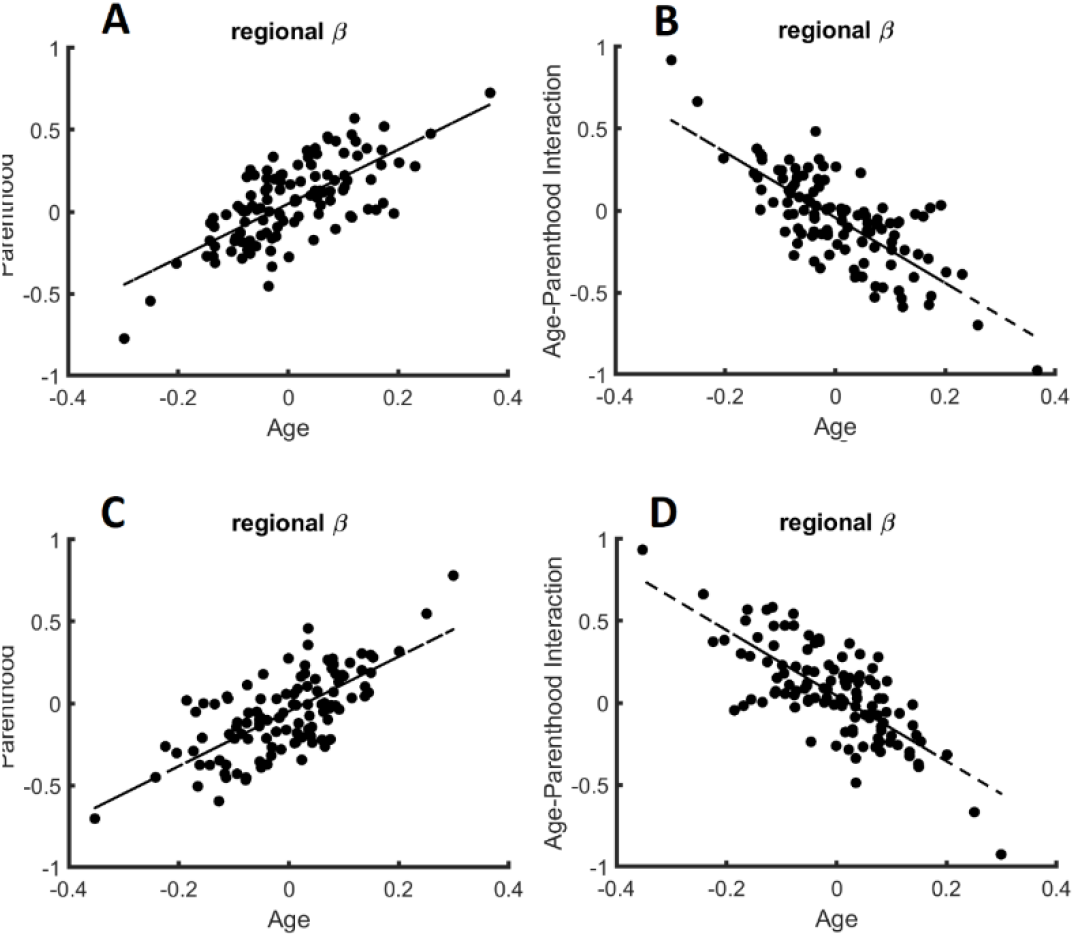
The relation of regional age and parenthood beta values in the linear models that predict whole-brain controllability in all subjects pooled together. Similar to the whole-brain models of controllability, the beta values for age and parenthood have the same signs for models of average (A) and modal (C) controllability while age and age-parenthood have opposite signs in models of (B) average and (D) modal controllability.

**Figure S4:**
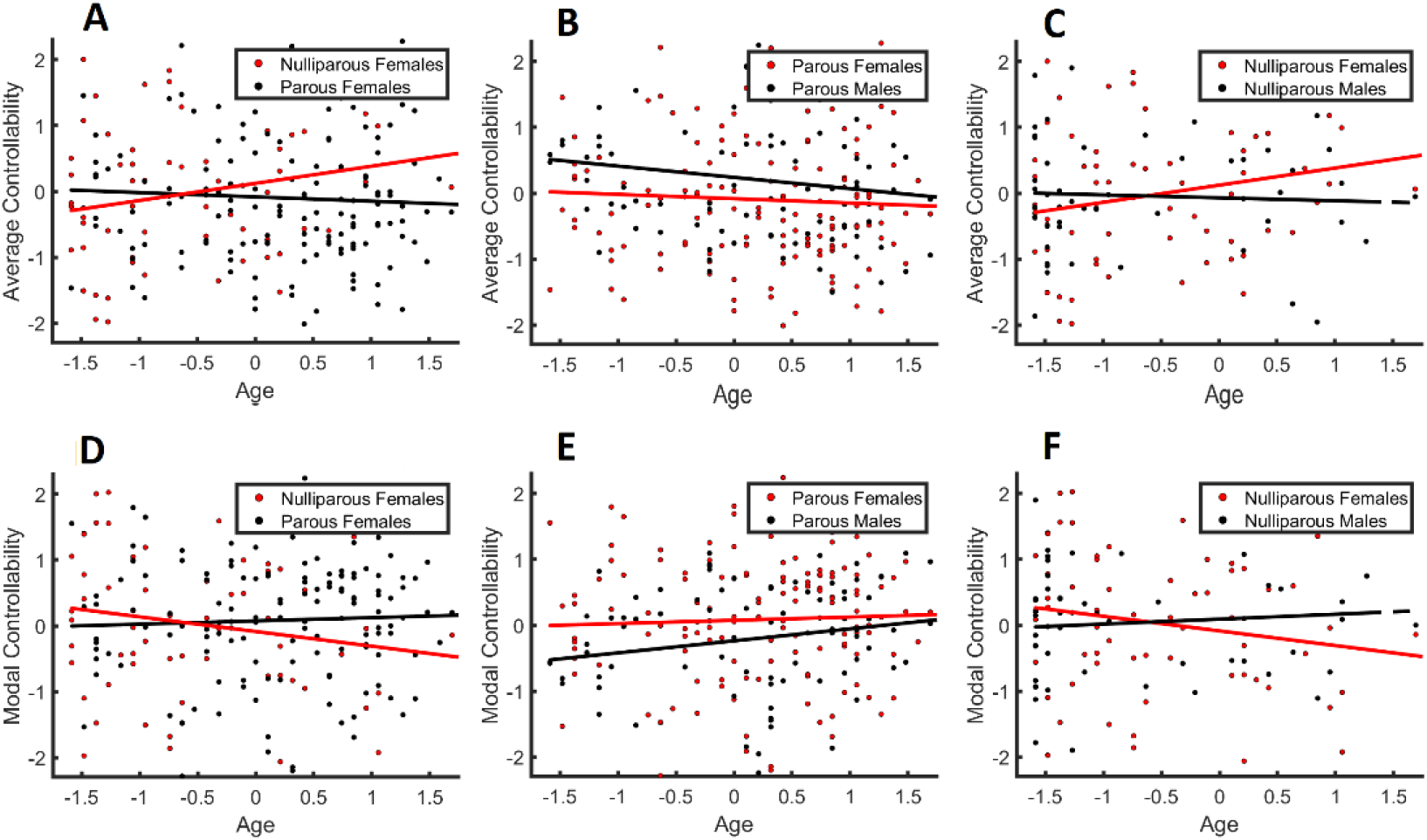
The relation between age, sex, and controllability together with the raw data. (A, D) Parenthood effect: average and modal controllability changes with age only in nulliparous females. (B, E) Parenthood by sex interaction: In parents, average controllability decreases with age and modal controllability increases although the effect is larger in mothers. (C, F) Parenthood by sex interaction: In non-parents, the age-related changes to controllability are more evident in nulliparous females compared to males.

**Table S1:** Linear model to estimate whole-brain average controllability for nulliparous women

| **Model: average controllability ~ 1 + age + site + structural strength** |  |  |  |  |
| --- | --- | --- | --- | --- |
|  | **Estimate** | **P-Value** | **T-value** | **SE** |
| **Intercept** | 8.3e-14 | 1 | 8e-13 | 0.11 |
| **Age** | 0.29 | 0.01 | 2.55 | 0.11 |
| **Site 1** | 0.16 | 0.26 | 1.13 | 0.15 |
| **Site 2** | 0.21 | 0.15 | 1.45 | 0.15 |
| **Structural Strength** | 0.37 | 2e-3 | 3.26 | 0.11 |
| Number of observations: 67, Error degrees of freedom: 62<br>Root Mean Squared Error: 0.901<br>R-squared: 0.238, Adjusted R-Squared: 0.189<br>F-statistic vs. constant model: 4.85, p-value = 0.00182 |  |  |  |  |

**Table S2:** Linear model to estimate whole-brain modal controllability for nulliparous women

| **Model: modal controllability ~ 1 + age + site + structural strength** |  |  |  |  |
| --- | --- | --- | --- | --- |
|  | **Estimate** | **P-Value** | **T-value** | **SE** |
| **Intercept** | -3.2e-14 | 1 | -2.8e-13 | 0.11 |
| **Age** | -0.26 | 0.03 | -2.21 | 0.12 |
| **Site 1** | -0.14 | 0.33 | -0.97 | 0.15 |
| **Site 2** | -0.22 | 0.14 | -1.47 | 0.15 |
| **Structural Strength** | -0.35 | 3.7e-3 | -3.01 | 0.12 |
| Number of observations: 67, Error degrees of freedom: 62<br>Root Mean Squared Error: 0.918<br>R-squared: 0.209, Adjusted R-Squared: 0.158<br>F-statistic vs. constant model: 4.09, p-value = 0.00525 |  |  |  |  |

**Table S3:** Linear model to estimate whole-brain average controllability for parous women

| **Model: average controllability ~ 1 + age + site + structural strength** |  |  |  |  |
| --- | --- | --- | --- | --- |
|  | **Estimate** | **P-Value** | **T-value** | **SE** |
| **Intercept** | 4.2e-14 | 1 | -1.9e-12 | 0.07 |
| **Age** | -0.05 | 0.51 | -0.66 | 0.07 |
| **Site 1** | 0.07 | 0.46 | 0.74 | 0.09 |
| **Site 2** | 0.18 | 0.05 | 2.00 | 0.09 |
| **Structural Strength** | 0.40 | 1.3e-7 | 5.54 | 0.07 |
| Number of observations: 157, Error degrees of freedom: 152<br>Root Mean Squared Error: 0.91<br>R-squared: 0.193, Adjusted R-Squared: 0.172<br>F-statistic vs. constant model: 9.09, p-value = 1.32e-06 |  |  |  |  |

**Table S4:** Linear model to estimate whole-brain modal controllability for parous women

| **Model: average controllability ~ 1 + age + site + structural strength** |  |  |  |  |
| --- | --- | --- | --- | --- |
|  | **Estimate** | **P-Value** | **T-value** | **SE** |
| **Intercept** | -3.7e-15 | 1 | -5.06e-12 | 0.07 |
| **Age** | 0.04 | 0.62 | 0.49 | 0.07 |
| **Site 1** | -0.08 | 0.4 | -0.83 | 0.09 |
| **Site 2** | -0.19 | 0.04 | -2.01 | 0.09 |
| **Structural Strength** | -0.39 | 4.2e-7 | -5.29 | 0.07 |
| Number of observations: 157, Error degrees of freedom: 152 |  |  |  |  |
| Root Mean Squared Error: 0.917 |  |  |  |  |
| R-squared: 0.18, Adjusted R-Squared: 0.158 |  |  |  |  |
| F-statistic vs. constant model: 8.33, p-value = 4.21e-06 |  |  |  |  |

**Table S5:**
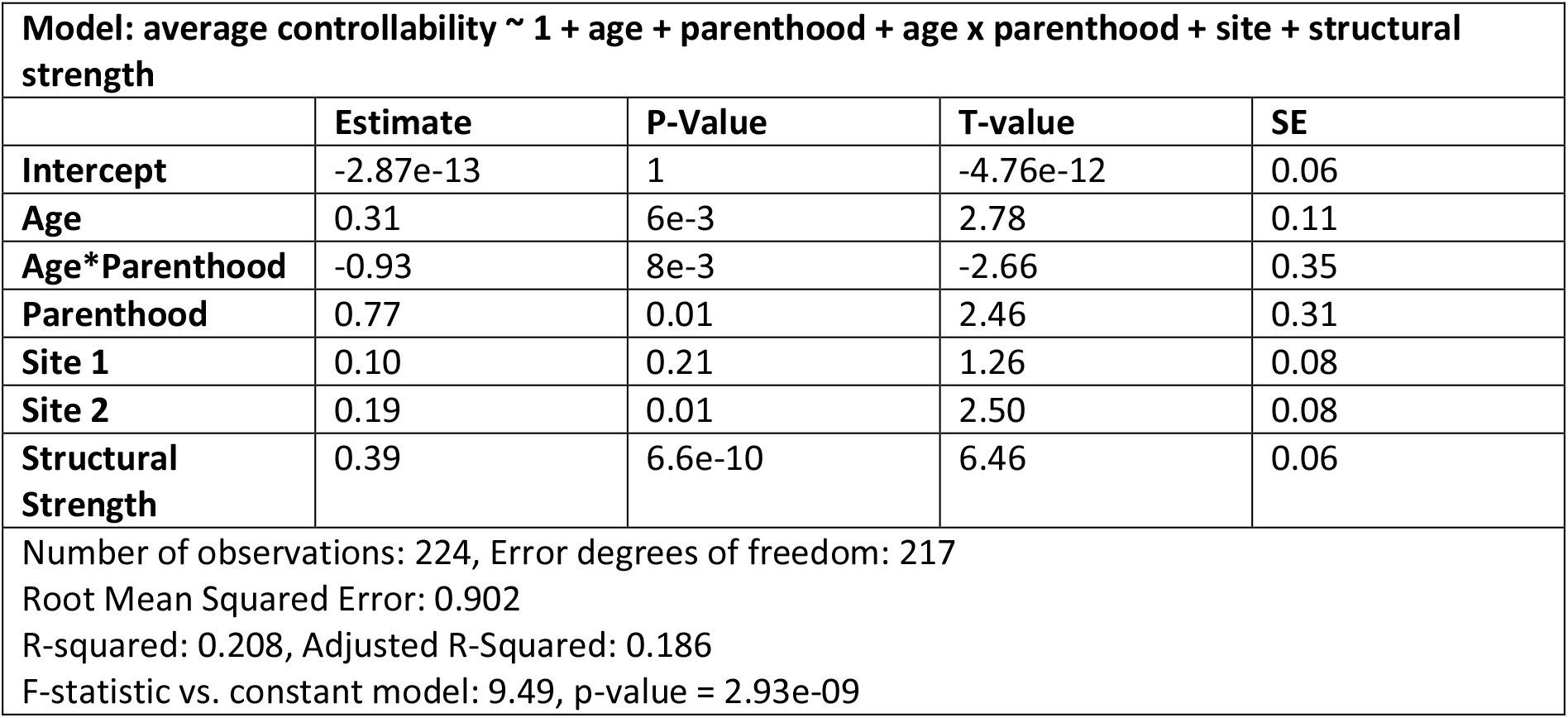
Full linear model to estimate whole-brain average controllability for women

| **Model: average controllability ~ 1 + age + parenthood + age x parenthood + site + structural strength** |  |  |  |  |
| --- | --- | --- | --- | --- |
|  | **Estimate** | **P-Value** | **T-value** | **SE** |
| **Intercept** | -2.87e-13 | 1 | -4.76e-12 | 0.06 |
| **Age** | 0.31 | 6e-3 | 2.78 | 0.11 |
| **Age\*Parenthood** | -0.93 | 8e-3 | -2.66 | 0.35 |
| **Parenthood** | 0.77 | 0.01 | 2.46 | 0.31 |
| **Site 1** | 0.10 | 0.21 | 1.26 | 0.08 |
| **Site 2** | 0.19 | 0.01 | 2.50 | 0.08 |
| **Structural Strength** | 0.39 | 6.6e-10 | 6.46 | 0.06 |
| Number of observations: 224, Error degrees of freedom: 217 |  |  |  |  |
| Root Mean Squared Error: 0.902 |  |  |  |  |
| R-squared: 0.208, Adjusted R-Squared: 0.186 |  |  |  |  |
| F-statistic vs. constant model: 9.49, p-value = 2.93e-09 |  |  |  |  |

**Table S6:** Full linear model to estimate whole-brain modal controllability for women

| **Model: average controllability ~ 1 + age + parenthood + age x parenthood + site + structural strength** |  |  |  |  |
| --- | --- | --- | --- | --- |
|  | **Estimate** | **P-Value** | **T-value** | **SE** |
| **Intercept** | -2.58e-13 | 1 | -4.23e-12 | 0.06 |
| **Age** | -0.27 | 0.02 | -2.40 | 0.11 |
| **Age\*Parenthood** | 0.80 | 0.02 | 2.25 | 0.35 |
| **Parenthood** | -0.66 | 0.04 | -2.11 | 0.31 |
| **Site 1** | -0.10 | 0.21 | -1.24 | 0.08 |
| **Site 2** | -0.20 | 0.01 | -2.52 | 0.08 |
| **Structural Strength** | -0.38 | 4.1e-9 | -6.13 | 0.06 |
| Number of observations: 224, Error degrees of freedom: 217 |  |  |  |  |
| Root Mean Squared Error: 0.913 |  |  |  |  |
| R-squared: 0.189, Adjusted R-Squared: 0.167 |  |  |  |  |
| F-statistic vs. constant model: 8.45, p-value = 2.98e-08 |  |  |  |  |

**Table S7:**
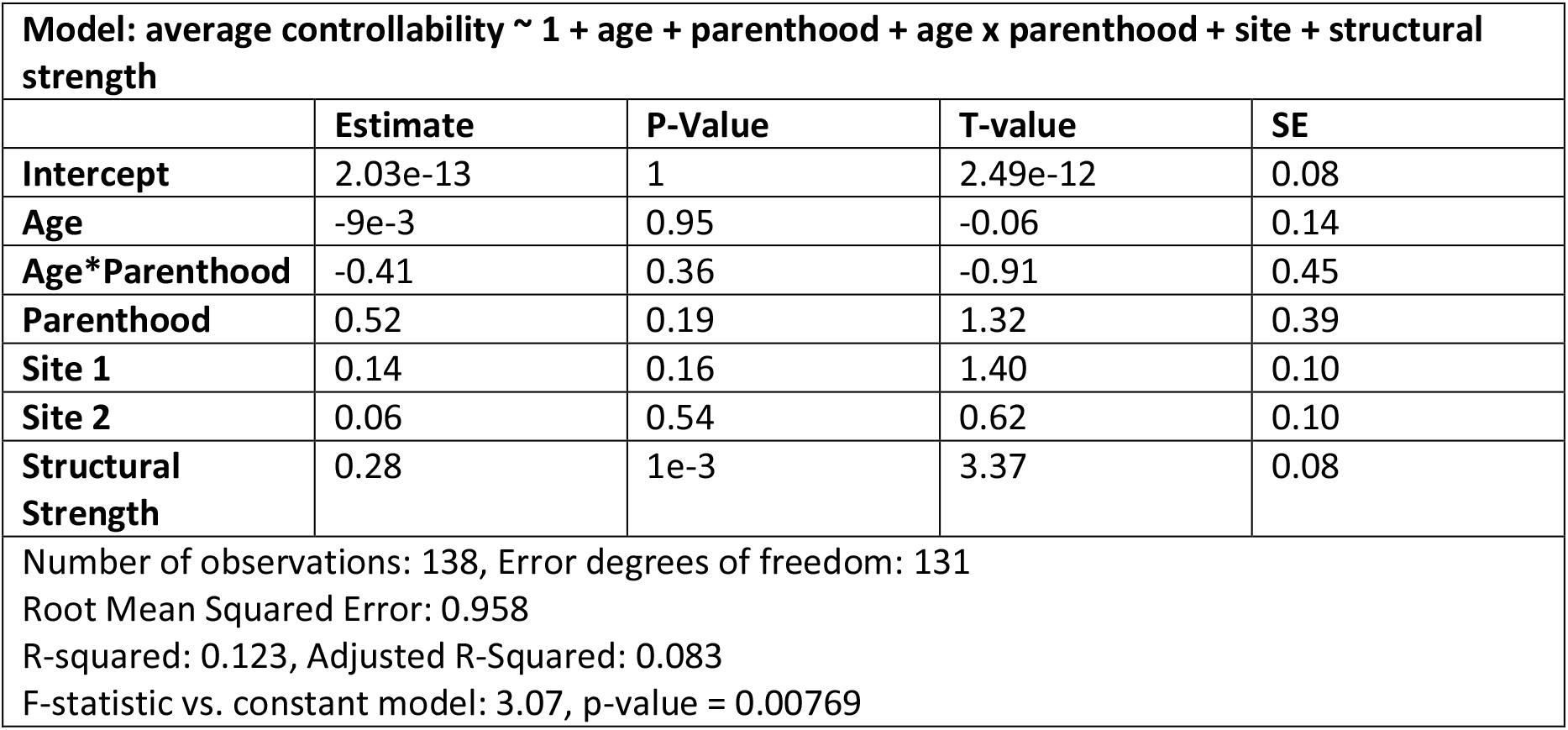
Full linear model to estimate whole-brain average controllability for men

| **Model: average controllability ~ 1 + age + parenthood + age x parenthood + site + structural strength** |  |  |  |  |
| --- | --- | --- | --- | --- |
|  | **Estimate** | **P-Value** | **T-value** | **SE** |
| **Intercept** | 2.03e-13 | 1 | 2.49e-12 | 0.08 |
| **Age** | -9e-3 | 0.95 | -0.06 | 0.14 |
| **Age\*Parenthood** | -0.41 | 0.36 | -0.91 | 0.45 |
| **Parenthood** | 0.52 | 0.19 | 1.32 | 0.39 |
| **Site 1** | 0.14 | 0.16 | 1.40 | 0.10 |
| **Site 2** | 0.06 | 0.54 | 0.62 | 0.10 |
| **Structural Strength** | 0.28 | 1e-3 | 3.37 | 0.08 |
| Number of observations: 138, Error degrees of freedom: 131<br>Root Mean Squared Error: 0.958<br>R-squared: 0.123, Adjusted R-Squared: 0.083<br>F-statistic vs. constant model: 3.07, p-value = 0.00769 |  |  |  |  |

**Table S8:** Full linear model to estimate whole-brain modal controllability for men

| **Model: average controllability ~ 1 + age + parenthood + age x parenthood + site + structural strength** |  |  |  |  |
| --- | --- | --- | --- | --- |
|  | **Estimate** | **P-Value** | **T-value** | **SE** |
| **Intercept** | 3.25e-13 | 1 | 3.97e-12 | 0.08 |
| **Age** | 0.04 | 0.77 | 0.29 | 0.14 |
| **Age\*Parenthood** | 0.35 | 0.44 | 0.78 | 0.45 |
| **Parenthood** | -0.48 | 0.22 | -1.22 | 0.40 |
| **Site 1** | -0.12 | 0.21 | -1.27 | 0.10 |
| **Site 2** | -0.02 | 0.84 | -0.20 | 0.10 |
| **Structural Strength** | -0.25 | 3e-3 | -3.00 | 0.08 |
| Number of observations: 138, Error degrees of freedom: 131<br>Root Mean Squared Error: 0.964<br>R-squared: 0.111, Adjusted R-Squared: 0.0703<br>F-statistic vs. constant model: 2.73, p-value = 0.0158 |  |  |  |  |

**Table S9:** Full linear model to estimate whole-brain average controllability for women and men

| **Model: average controllability ~ 1 + age + parenthood + age x parenthood + site + structural strength** |  |  |  |  |
| --- | --- | --- | --- | --- |
|  | **Estimate** | **P-Value** | **T-value** | **SE** |
| **Intercept** | -2.32e-14 | 1 | -4.79e-13 | 0.05 |
| **Age** | 0.17 | 0.04 | 2.02 | 0.09 |
| **Age\*Parenthood** | -0.70 | 0.01 | -2.57 | 0.27 |
| **Parenthood** | 0.63 | 9e-3 | 2.62 | 0.24 |
| **Site 1** | 0.11 | 0.06 | 1.91 | 0.06 |
| **Site 2** | 0.14 | 0.02 | 2.37 | 0.06 |
| **Structural Strength** | 0.37 | 3.01e-13 | 7.58 | 0.05 |
| Number of observations: 362, Error degrees of freedom: 355<br>Root Mean Squared Error: 0.921<br>R-squared: 0.166, Adjusted R-Squared: 0.152<br>F-statistic vs. constant model: 11.8, p-value = 4.7e-12 |  |  |  |  |

**Table S10:**
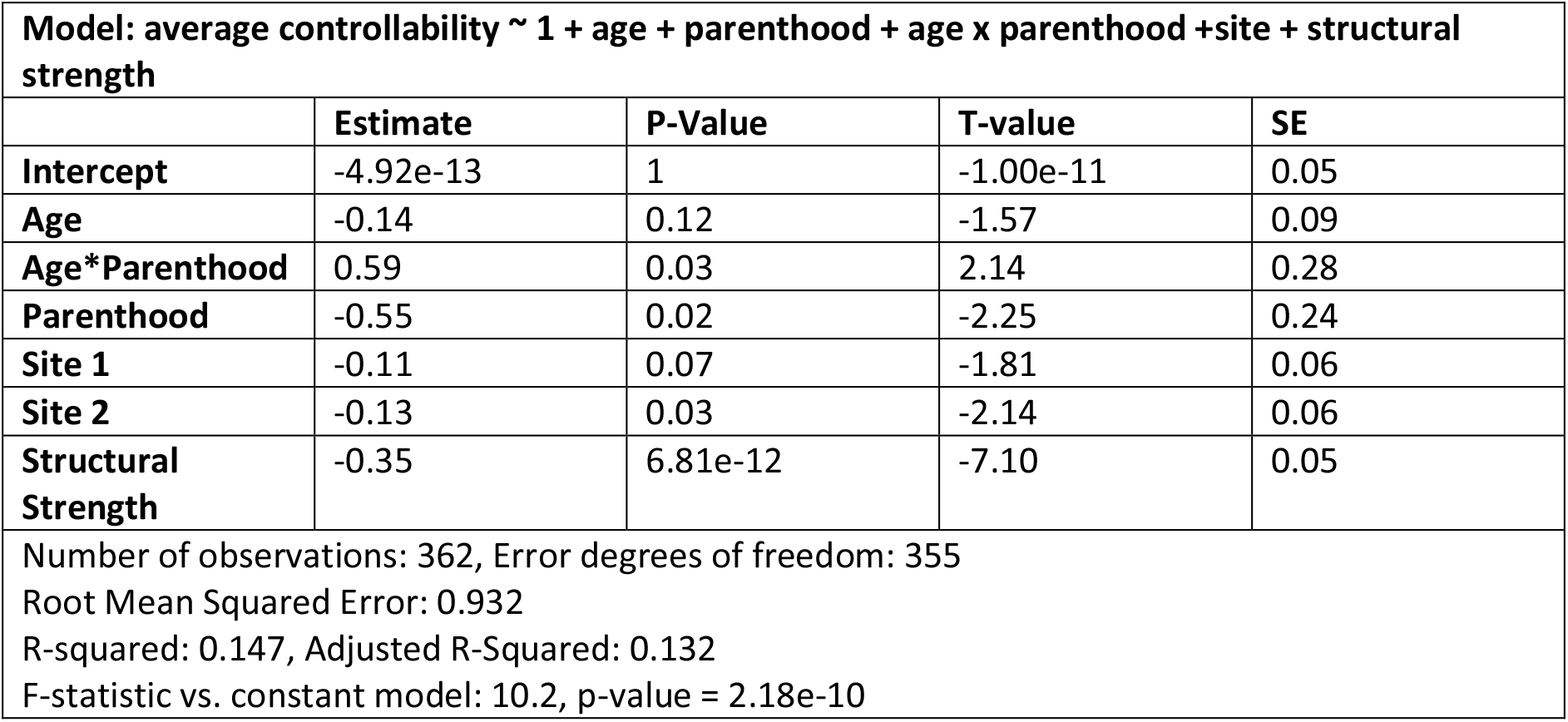
Full linear model to estimate whole-brain modal controllability for women and men

**Table S11:** Full linear model to estimate whole-brain average controllability for women and men with gender as an additive covariate

| **Model: average controllability ~ 1 + age + parenthood + age x parenthood +site + structural strength** |  |  |  |  |
| --- | --- | --- | --- | --- |
|  | **Estimate** | **P-Value** | **T-value** | **SE** |
| **Intercept** | -2.43e-14 | 1 | -5.03e-13 | 0.05 |
| **Age** | 0.18 | 0.04 | 2.05 | 0.09 |
| **Age\*Parenthood** | -0.71 | 0.01 | -2.58 | 0.27 |
| **Parenthood** | 0.64 | 8e-3 | 2.64 | 0.24 |
| **Site 1** | 0.12 | 0.05 | 1.94 | 0.06 |
| **Site 2** | 0.14 | 0.02 | 2.40 | 0.06 |
| **Structural Strength** | 0.36 | 1.53e-12 | 7.33 | 0.05 |
| **Gender** | -0.03 | 0.48 | -0.71 | 0.05 |
| Number of observations: 362, Error degrees of freedom: 354<br>Root Mean Squared Error: 0.922<br>R-squared: 0.167, Adjusted R-Squared: 0.151<br>F-statistic vs. constant model: 10.2, p-value = 1.35e-11 |  |  |  |  |

**Table S12:**
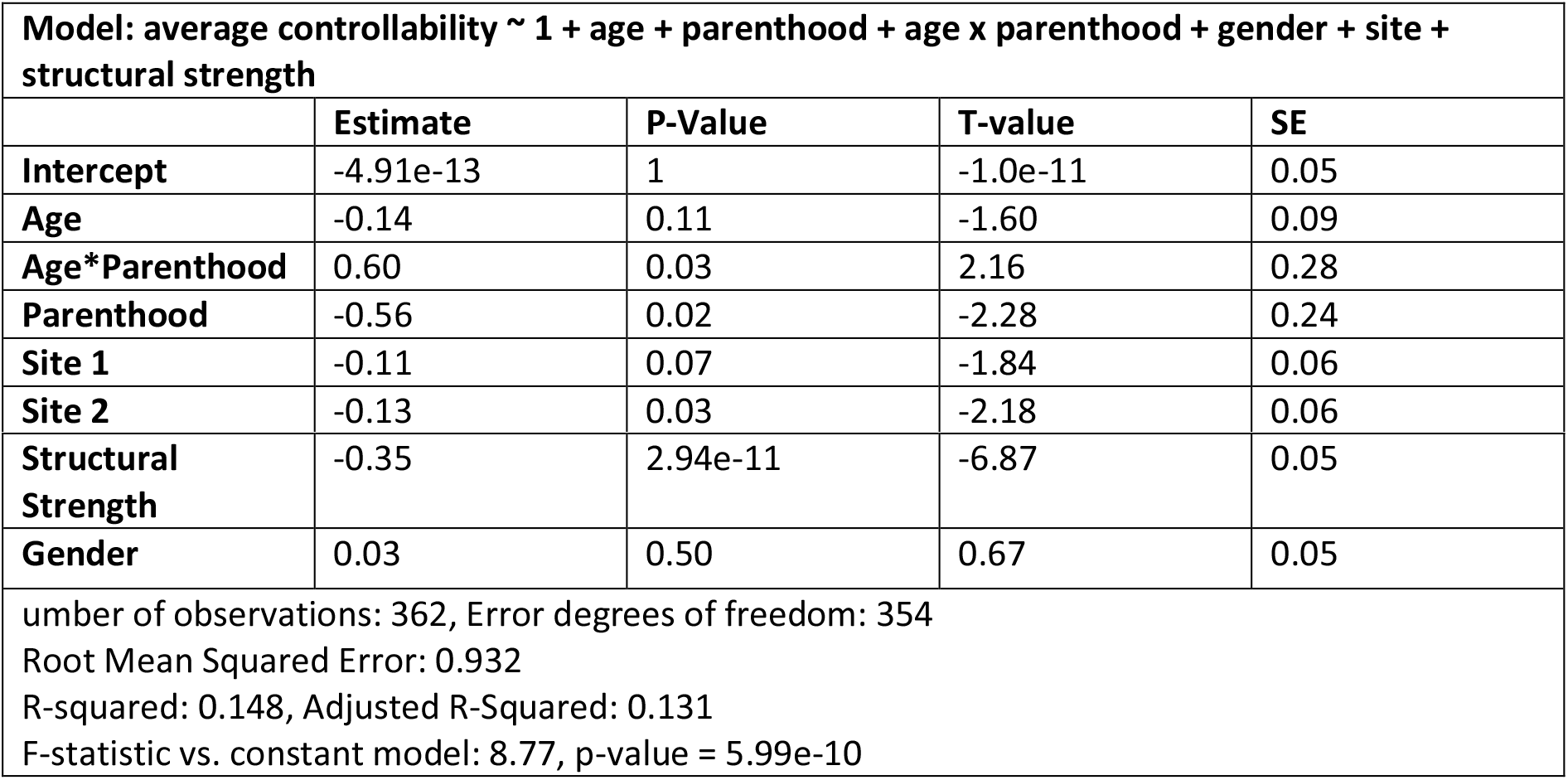
Full linear model to estimate whole-brain modal controllability for women and men with gender as an additive covariate

**Table S13:** Replication analysis of linear models with different age intervals. For each model, we replicated the results for data with age > age-limit where the age-limit was increased from 18 to 50 years old.

| | $\beta_{age}$ with age > 30 | $\beta_{age}$ with age between [18 50] years old |
| --- | --- | --- |
| **Table S1** | 0.29 | 0.31±0.14 |
| **Table S2** | -0.26 | -0.28±0.14 |
| **Table S3** | -0.05 | -0.06±0.03 |
| **Table S4** | 0.04 | 0.03±0.03 |
| **Table S5** | 0.31 | 0.27±0.09 |
| **Table S6** | -0.27 | -0.25±0.09 |
| **Table S7** | -9e-3 | -0.02±0.11 |
| **Table S8** | 0.04 | 0.03±0.10 |
| **Table S9** | 0.17 | 0.19±0.06 |
| **Table S10** | -0.14 | -0.17±0.07 |
| **Table S11** | 0.18 | 0.19±0.06 |
| **Table S12** | -0.14 | -0.17±0.07 |

